# Human microbiome aging clocks based on deep learning and tandem of permutation feature importance and accumulated local effects

**DOI:** 10.1101/507780

**Authors:** Fedor Galkin, Aleksandr Aliper, Evgeny Putin, Igor Kuznetsov, Vadim N. Gladyshev, Alex Zhavoronkov

## Abstract

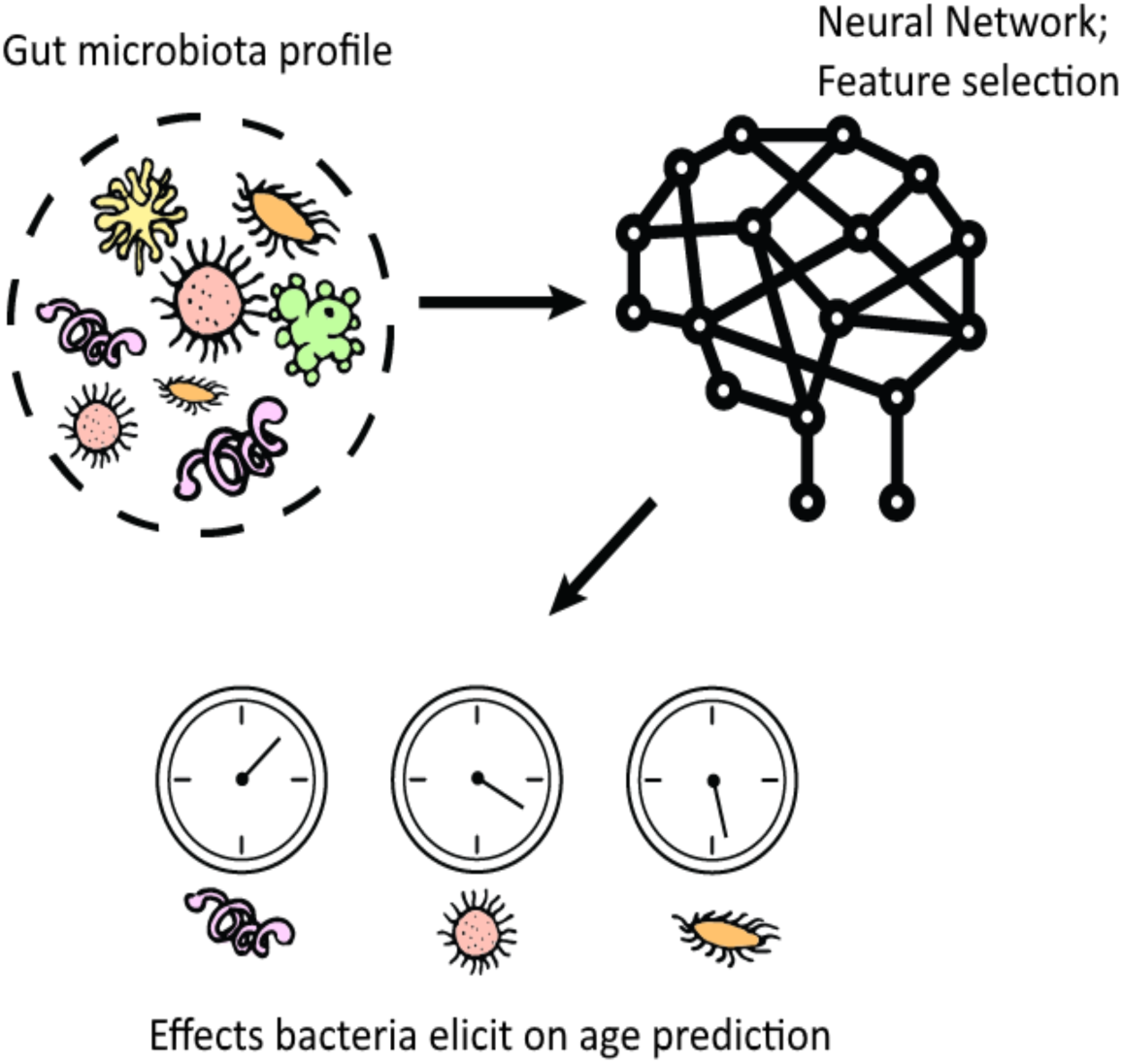

**Abstract:** The human gut microbiome is a complex ecosystem that both affects and is affected by its host status. Previous analyses of gut microflora revealed associations between specific microbes and host health and disease status, genotype and diet. Here, we developed a method of predicting biological age of the host based on the microbiological profiles of gut microbiota using a curated dataset of 1,165 healthy individuals (3,663 microbiome samples). Our predictive model, a human microbiome clock, has an architecture of a deep neural network and achieves the accuracy of 3.94 years mean absolute error in cross-validation. The performance of the deep microbiome clock was also evaluated on several additional populations. We further introduce a platform for biological interpretation of individual microbial features used in age models, which relies on permutation feature importance and accumulated local effects. This approach has allowed us to define two lists of 95 intestinal biomarkers of human aging. We further show that this list can be reduced to 39 taxa that convey the most information on their host’s aging. Overall, we show that (a) microbiological profiles can be used to predict human age; and (b) microbial features selected by models are age-related.

## Introduction

The human gut is colonized by a dense microbial community, calculated to consist of 10^14^ cells, which is an order of magnitude higher than the number of cells in the host ^1^. Gut microbiota is a complex ecosystem that carries multiple important functions in the organism. Apart from being a core element of the digestive system, microbiota regulates immunity, processes xenobiotics, produces important metabolites, and even affects higher neural functions ^2–4^. The influence, however, is not one-sided: microbiota is not simply determining certain host characteristics, as it responds to signals from the host via multiple feedback loops ^5^. Some of these feedback loops were found to be reflected in the microbiota composition.

For example, multiple studies indicate that irritable bowel diseases can develop following the intense immune response to an intestinal infection. Microbiota responds to proinflammatory milieu with a decreased number of beneficial bacteria that lack mechanisms to survive under such hostile conditions. In return, host immunity reacts to suppress the blooming pathogenic community, which produces chronic inflammation ^6^. Such changes constantly happen throughout an individual’s life and may be deleterious or beneficial, reflect strictly individual choices or be the effects of more widespread factors across populations.

Metagenomic studies have provided valuable insights into how the gut microflora progresses with age. They revealed that gut colonization occurs during birth with the bacteria living in the birth canal. The “pioneer microbiome” consists of facultative aerobes (e.g. *Escherichia, Enterococcus, etc.*) that gets replaced during breast feeding with obligate anaerobes (e.g. *Bifidobacterium infantis*) ^7^. Upon weaning, another community shift happens towards more adult-like microbiomes ^8^. These early stages of colonization are extremely important as normal infant microbiota promotes intestinal mucus formation, prevents pathogen blooming, and regulates T-cells. The importance of early colonization is further emphasized by studies that indicate higher occurrences of eczema and food allergies in children with atypical microbiota ^9^ development (e.g. increased abundance of *Clostridium* and *Escherichia* microbes) ^10^. Factors such as the mode of birth delivery (vaginal or cesarean), infant diet (breast milk or formula), and maternal microbiome greatly influence microbiome development.

Although infant microbiome succession is well studied and can be used to assess the risks of various health conditions, its transition to adult microbiome is less understood. More so, composition variability attributed to geographic location, medical history, diet, and other factors make it hard to analyze adult microbiomes as effectively as those of infants. Age-related studies of human microbiome have failed to produce a straightforward theory of gut flora aging. Some studies indicate decreasing biodiversity in the elderly gut ^11,12^. However, that is not the case for all data sets, and elderly healthy people may have microbiomes as diverse as the younger population ^13,14^. Other findings include changes in specific taxa abundance in aging microbiota. Such bacterial genera as *Bacteroides, Bifidobacterium, Blautia, Lactobacilli, Ruminococcus* have been shown to decrease in the elderly, while *Clostridium, Escherichia, Streptococci, Enterobacteria* increase ^15,16^. However, these patterns are not strictly established as results vary greatly across different studies. This may be attributed to different methodologies as well as unbalanced data sets that may contain people of different lifestyles ^17^.

Despite these complications, the consensus is that the elderly gut has lower counts of short chain fatty acid (SCFA) producers such as *Roseburia* and *Faecalibacterium* and an increased number of aerotolerant and pathogenic bacteria. Such shifts can lead to dysbiosis, which in turn contributes to the onset of multiple age-related diseases ^9^. The idea that the gut microflora can be a major contributor to the aging process is not new. Already in the beginning of the 20^th^ century, a Nobel Prize-winning Russian scientist Ilya Metchnikoff proposed that the malicious microbes processing undigested food (especially peptolytic bacteria, e.g. *Escherichia* and *Clostridium*) lead to autointoxication. Treating autointoxication with pro- and pre-biotics (such as *Lactobacillus* preparations) was suggested to alleviate an age-associated decline in organismal function. Recent studies have demonstrated promising results in line with this century-old hypothesis ^18–20^.

The standard way of separating the gut microbiome into three chronological states - child, adult, and elderly microbiomes - lack a clear set of rules. Among them, adult microbiome remains the greatest mystery. It has no established succession stages, as in newborns, and does not normally reflect gradient detrimental processes typical for an old organism. This poses a question whether normal adult microbiome progresses at all or it is in a state of stasis. Considering the aging process is gradual and involves accumulation of damage and other deleterious changes ^21^ (as also indicated by a number of biomarkers such as DNA methylation clocks ^22,23^), it is logical to suppose that gut microbiome succession is also gradual ^24^. However, attempts to use microbiome-derived features to predict chronological age have been inconclusive. A support vector machine model trained on human metagenomic data to classify samples as young or old was shown to be only 10-15% more accurate than random assignment, as indicated by the Area Under the Curve (AUC) score ^25^. Another study attempting to use a co-abundance clustering approach has demonstrated general trendlines of microbiota composition for hosts aged 0-100 ^26^. According to the study, specific clades of the gut community significantly differ in abundance among young adults compared to the middle aged. However, the lack of dietary and lifestyle data prevents the authors from putting together a conclusive theory of gut microflora progression. Compared to the well-established DNAm aging clocks that achieve mean absolute error (MAE) <5 years, these results of microflora-based age prediction suggest much room for improvement ^27,28^.

The renaissance of deep learning that started in 2015 resulted in unprecedented machine learning performance in image, voice, and text recognition, as well as a range of biomedical applications ^29^ such as drug repurposing ^30^ and target identification ^31^. One of the most impactful applications of DL in biomedicine was in the applications of generative models to *de novo* molecular design ^32–36^. In the context of aging research, these new methods can be combined for geroprotector discovery ^37–41^. Indeed, since 2013, many aging clocks have been developed in both humans and other model organisms. The published aging clocks utilizing deep learning were developed using standard clinical blood tests ^42^, facial images ^43^, physical activity data, ^44^ and transcriptomic data ^45^. These clocks were used to rank the most important features contributing to the accuracy of the prediction by using the permutation feature importance (PFI), deep feature selection (DFS) and other techniques. These clocks were also used to assess the population-specificity of the various data types ^42^.

The goal of this study was to build a predictor of age with whole genome sequencing (WGS) data aggregated from multiple sources and various machine learning techniques and use it to examine patterns of incessant microflora succession. Here, we report a method to estimate a host’s age based on their microflora taxonomic profile, assess the importance of specific taxa in organismal aging, and suggest candidate geroprotective microbiological interventions.

## Methods

### Data acquisition

Only publicly available, fully anonymized data sets from WGS human metagenomic studies deposited in ENA and SRA were used. The corresponding project IDs are: <u>ERP005534</u>, <u>SRP008047</u>, <u>ERP009422</u>, <u>ERP004605</u>, <u>ERP002061</u>, <u>ERP002469</u>, <u>ERP019502</u>, <u>SRP002163</u>, <u>ERP003612</u>, <u>ERP008729</u>.^46–49^ Only healthy individuals with age metadata available were included in this study. These individuals were from Austria, China, Denmark, France, Germany, Kazakhstan, Spain, Sweden and USA. aged 20-90 years old. In total, 1,165 healthy individuals and 3,663 samples from 10 publicly available datasets were aggregated and analyzed (Figure 1).

**Figure 1:**
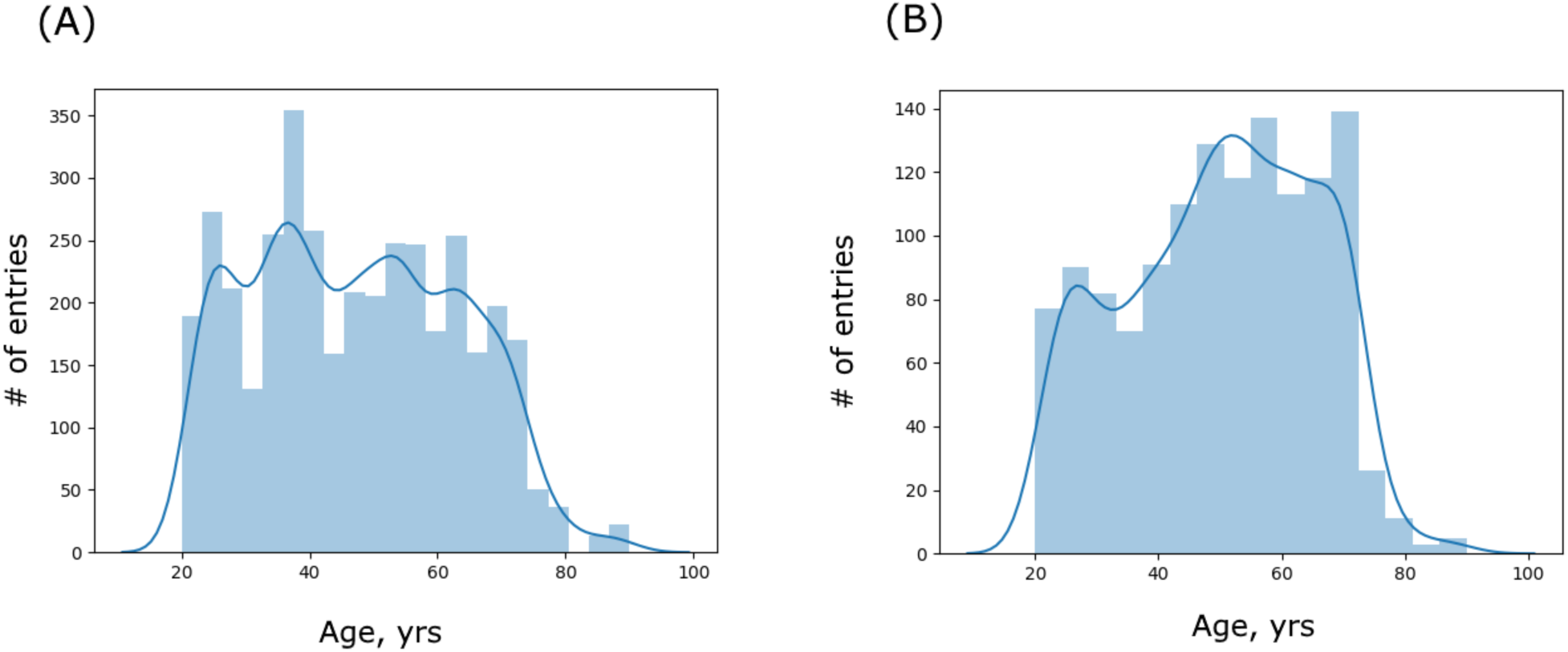
age distributions for 3’663 runs (A) and 1’165 donors (B) used in this study.

### Abundance calculation

All acquired sequencing files have been quality trimmed and quality filtered with <u>BBTools</u>^50^. Human sequences have been detected using hg19 genome index. Additionally, specimen dilution test has been carried out as specified in ^51^. Resulting reads have been analyzed with <u>Centrifuge</u> and mapped against the collection of bacterial and archaeal genomes ^52^. In certain cases, operational taxonomic units ables have been modified to exclude unreliably detected microbes (relative abundance < 1e-5) and minor microbial species (<1.3e-3 prevalence). No sample has lost more than 5% of its abundance. After all the modifications, individual taxonomic profiles have been renormalized by dividing the vector by the sum of the abundances left.

### Neural networks training

#### Regression

All deep neural networks (DNNs) were implemented using the Python 3.6 <u>Keras</u> library with <u>Tensorflow</u> backend. Feature selection models were trained using a full list of species-level features, which includes 1,673 microbial taxa. Training and validation sets were separated to contain 90% and 10% of all profiles in all cases. Two regressors were built: one using taxonomic profiles derived from individual samples (sample-based model) and a second one using taxonomic profiles averaged among all the samples belonging to the same host (host-based model). Models were trained as a regressor with five-fold cross-validation. After completing grid search for various model configurations, the best performing model was selected based on the maximal R^2^ score.

The best performing model architecture was determined in the sample-based setting. It contains three hidden layers with 512 nodes in each, with PReLU activation function, Adam optimizer, dropout fraction 0.5 at each layer, and 0.001 learning rate (Figure 2). The same architecture was applied to within the host-based setting. To verify the importance of features derived from the sample-based DNN model, gradient boosting was used, as implemented in <u>XGBoost</u> Python library ^53^. The best performing XGBoost model was trained using the following parameters: linear_nthread = 35, max_depth = 6, max_delta_step = 2, lambda= 0, gamma=0.1, eta=0.1, alpha = 0.5. The XGBoost models’ performance was evaluated using MAE.

**Figure 2:**
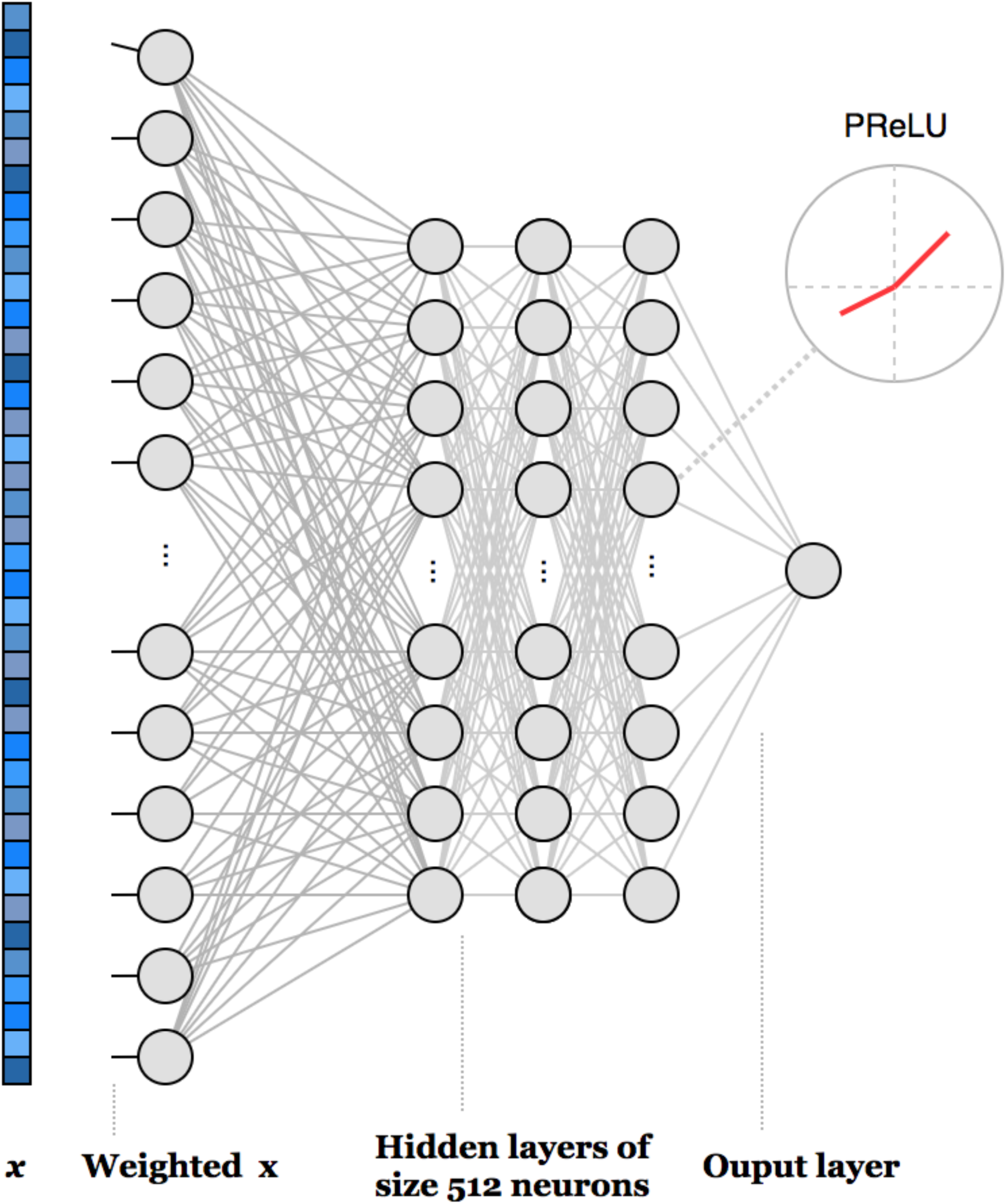
The neural network configuration for the best performing DNN regressor. The regressor takes in a full species level taxonomic profile and estimates the donor’s exact chronological age. The first hidden layer is linear and was used only to assess feature importance in accordance with deep feature selection method ^55^.

#### Classification

Age classifier models were trained using a subset of either 95 features or 39 features. Training and validation were separated to contain 80% and 20% of all donors, respectively. The age bracket classifier was implemented with the Python Keras library using Tensorflow backend. A weighted F1-score was selected as the target metric to assess model performance. Best performing architectures are illustrated in Figure 3. For 95 feature classifier it is: 128, 32 and 8 nodes respectively in 3 hidden layers, dropout rate of 0.5, PReLU activation function in hidden layers, softmax activation function in the output layer ^54^. For 39 feature classifier it is: [64, 8] nodes in 2 hidden layers, 0.5 dropout rate, PReLU activation function in hidden layers, Softmax activation in the output layer.

**Figure 3:**
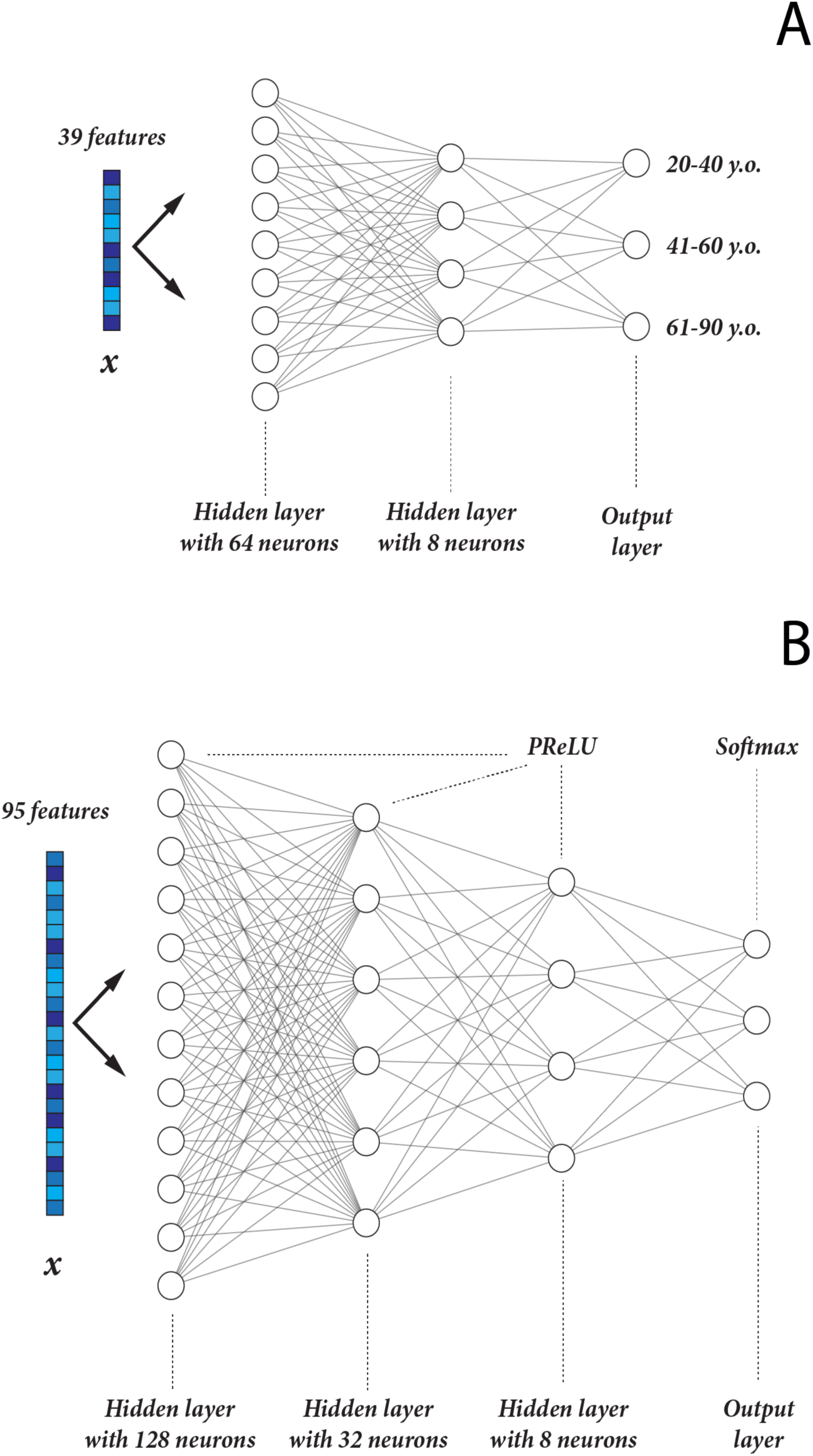
The best performing DNN configurations for age bracket classification (20-40, 41-60, 61-90 years) based on short marker sets: 39 taxa (A), 95 taxa (B).

### Oversampling

To solve the class imbalance problem while building models for age bracket prediction, we used oversampling. Self Organizing Maps (SOMs) based on presence/absence profiles (1 if a taxon is detected in a sample, 0 if it is not) have been built for each age bracket with the Python library <u>Somoclu</u>. Each SOM consists of 100 cells placed on a toroid lattice. To generate synthetic profiles for underrepresented classes, codebook vectors are picked at random with replacement according to the number of Best Matching Units (BMUs) mapped to them. Codebook values are used as probabilities for including a taxon into a fake sample. Fake presence/absence profiles are then multiplied by a vector of mean abundances of corresponding BMUs and normalized.

### Feature importance

To assess individual feature importance, we have applied the Permutation Feature Importance (PFI) technique. PFI measures the change in prediction quality (measured in R^2^ score decrease) upon permuting a single feature vector. Greater decrease in quality signal greater importance of the feature. The features deemed most important have been further assessed with the Accumulated Local Effects (ALE) method to determine the change in age prediction upon minor changes in a microbial species abundance. ALE has been implemented following the algorithm described below. For each of the 95 selected species, a quantile value table (with 5% steps) has been composed. Local Effects (LE) for each quantile bin have been calculated by measuring the average change in prediction upon substituting observed abundance of a feature, with right and left bin border values. ALEs for each quantile are calculated by adding up all the previous LEs and centering the result to make the average effect of each taxon zero.

## Results

### Age prediction using machine learning

To examine the relationship between human gut taxonomic profiles and chronological age, we prepared a collection of full metagenome sequences for 1,165 healthy individuals (3,663 samples total) from 10 publicly available datasets. All individuals in our data set were between 20 and 90 years, with median age of 46 years. After randomly separating the 3,663 samples into training (90%) and validation (10%) sets, we trained a deep neural network regressor to predict donor’s age using a vector of relative abundances for 1,673 microbial species. MAE achieved by the best model configuration was 3.94 years, with R^2^ of 0.81 (Figure 5A). We then divided the samples into three age groups (20-39, 40-59 and 60-90 years) and found that the predicted age distribution generated by the model closely matched the actual age distribution (Figure 6).

**Figure 4:**
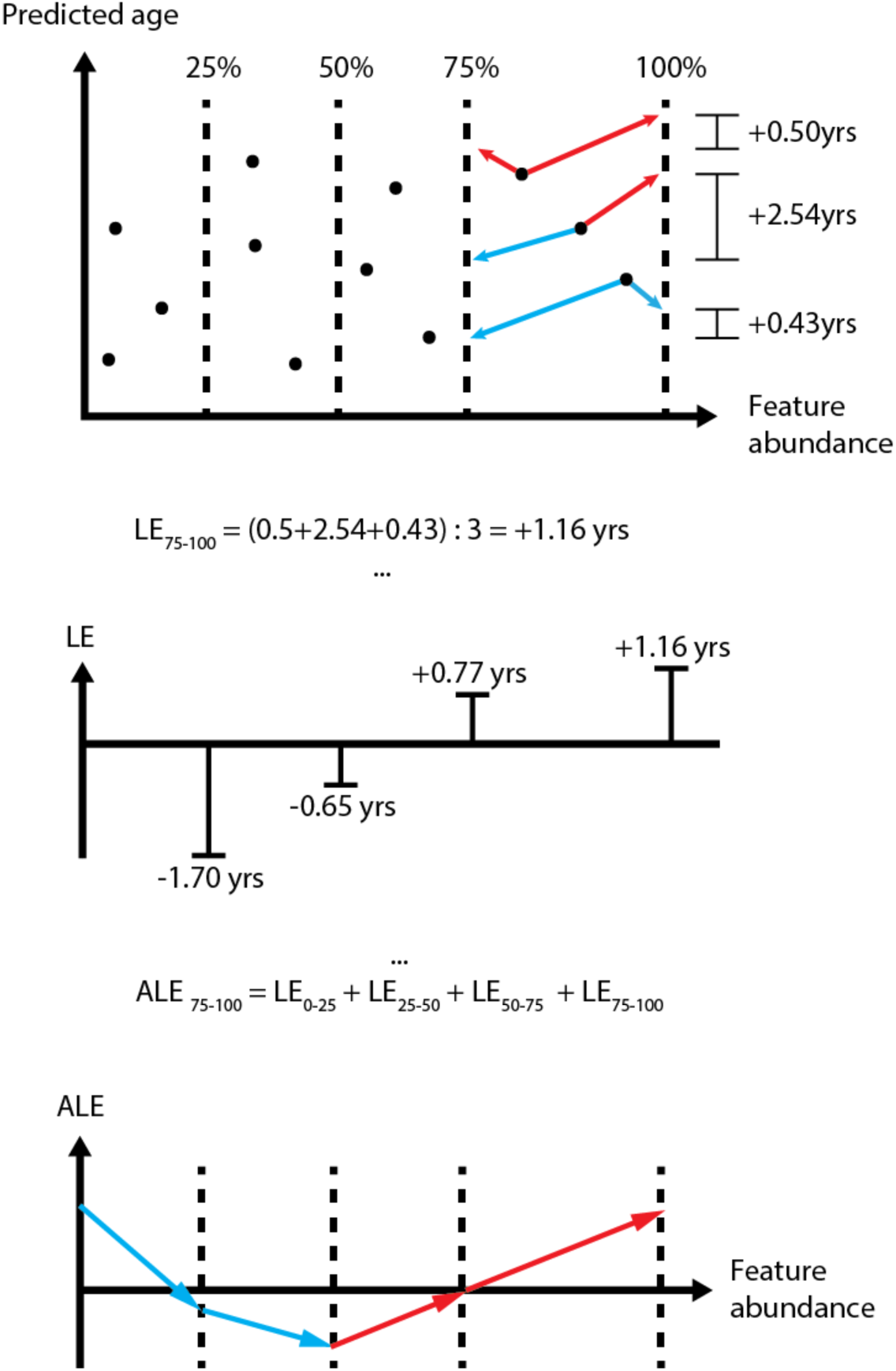
Accumulated Local Effects (ALE) method used in this paper to assess specific taxa influence on age prediction. Changes in predicted age upon substituting observed taxon abundance with quantile values are averaged and recorded for every quantile bin. Then, they are summed to produce ALEs, which are additionally centered for convenience.

**Figure 5:**
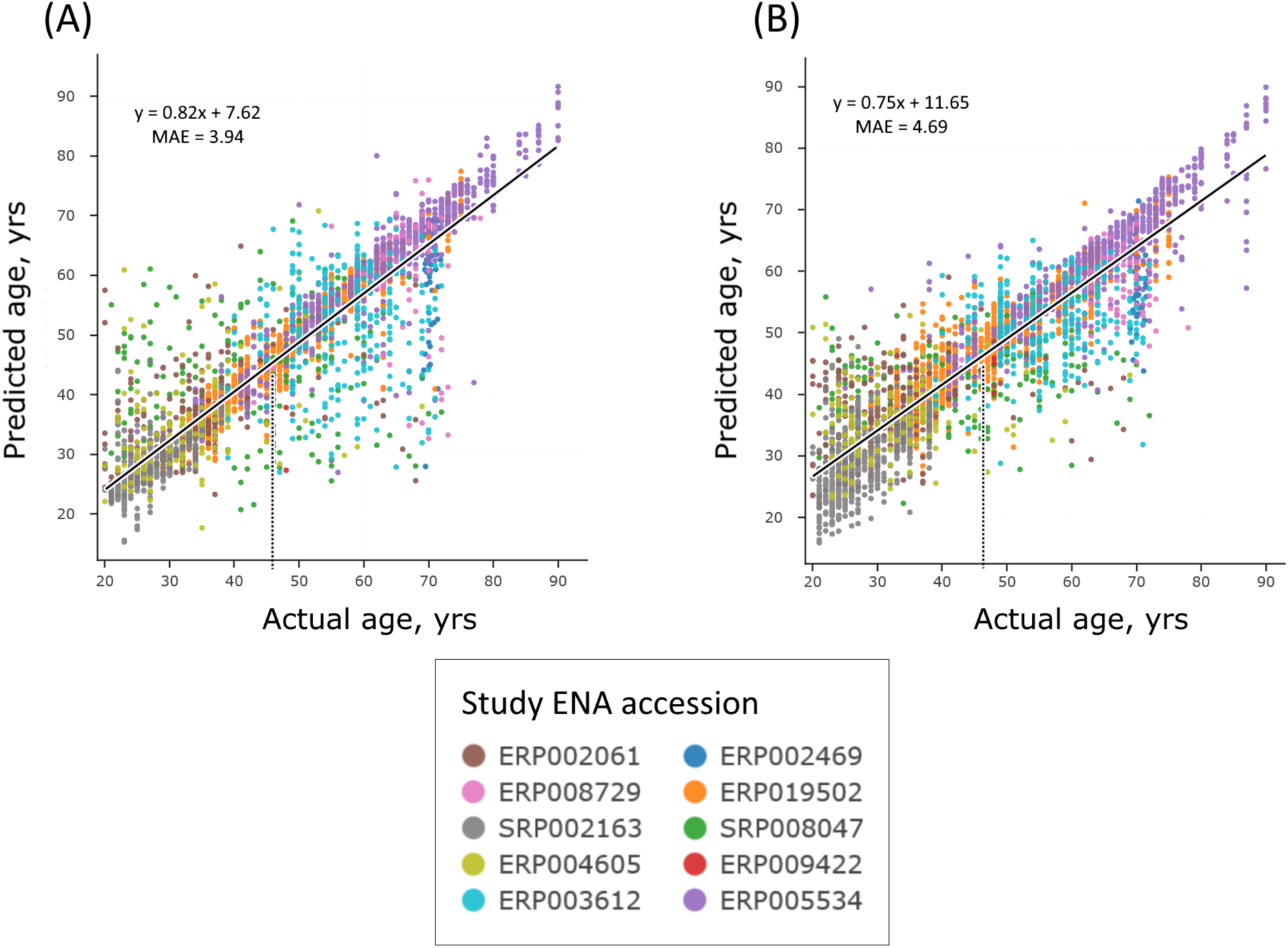
Age predictions derived from cross-validation of the sample-based DNN model (A) and the XGB model (B). Samples are colored by data source, and dashed lines mark the median of observed age (46 years).

**Figure 6:**
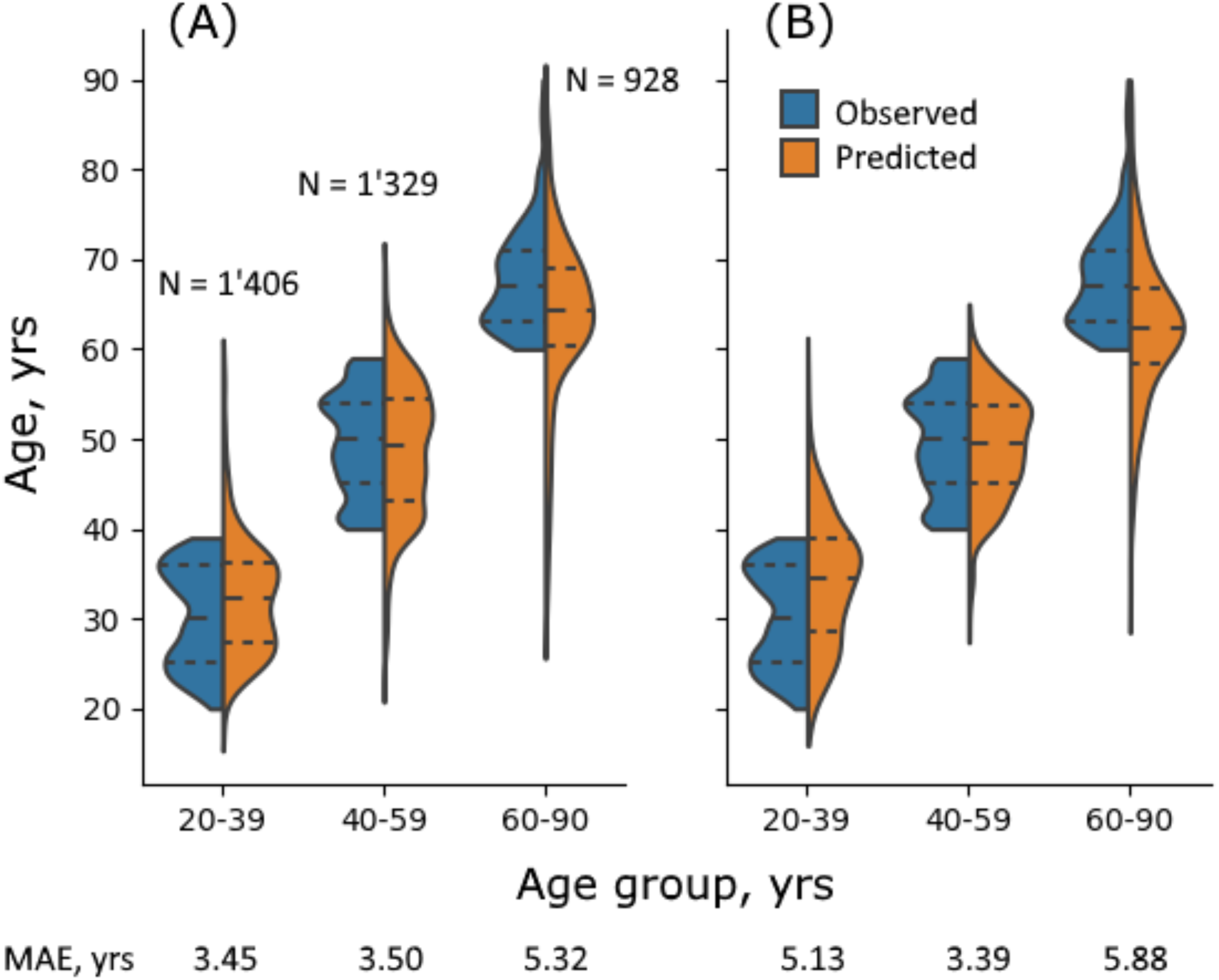
Density distribution for observed (blue) and predicted (orange) ages for two regressors: DNN (A) and XGB (B). “N” stands for the total number of samples per class. Dashed lines within violins stand for quantile borders. Mean Absolute Error (MAE) (in years) for each age group is marked below the graph.

To verify the results obtained with DNN, we implemented random forest, support vector machine and elastic net regressor. All of these methods performed poorly compared to the DNN approach with the mean absolute errors exceeding 11 years. Apart from them, we trained a gradient boosting (XGB) regressor with accuracy comparable to the DNN model (MAE = 4.69 years, R^2^ = 0.81) (Figure 5B). Both approaches skew the predictions towards the median age — 46 years (Figure 6). While there are certain variations within taxonomic profiles due to differences in geographical location or diet types, the described predictors can be applied to adult people from various populations equally well (see Supplementary).

### Microbiological influence on age prediction

Using Permutation Feature Importance (PFI), we assessed which taxa abundances play the greatest role in microbiological age prediction. We identified 95 features that decrease both XGB and DFS models’ R^2^ score by >0.001 (Figure 7). According to PFI scores, DNN regressor is more sensitive to highly abundant species, while XGB regressor contains some minor taxa among its most important features. We consider this an indication of DNN’s increased robustness compared to other methods. The complete list of 95 taxa with corresponding scores, abundances and prevalences can be found in <u>Supplementary Table 1</u>.

**Figure 7:**
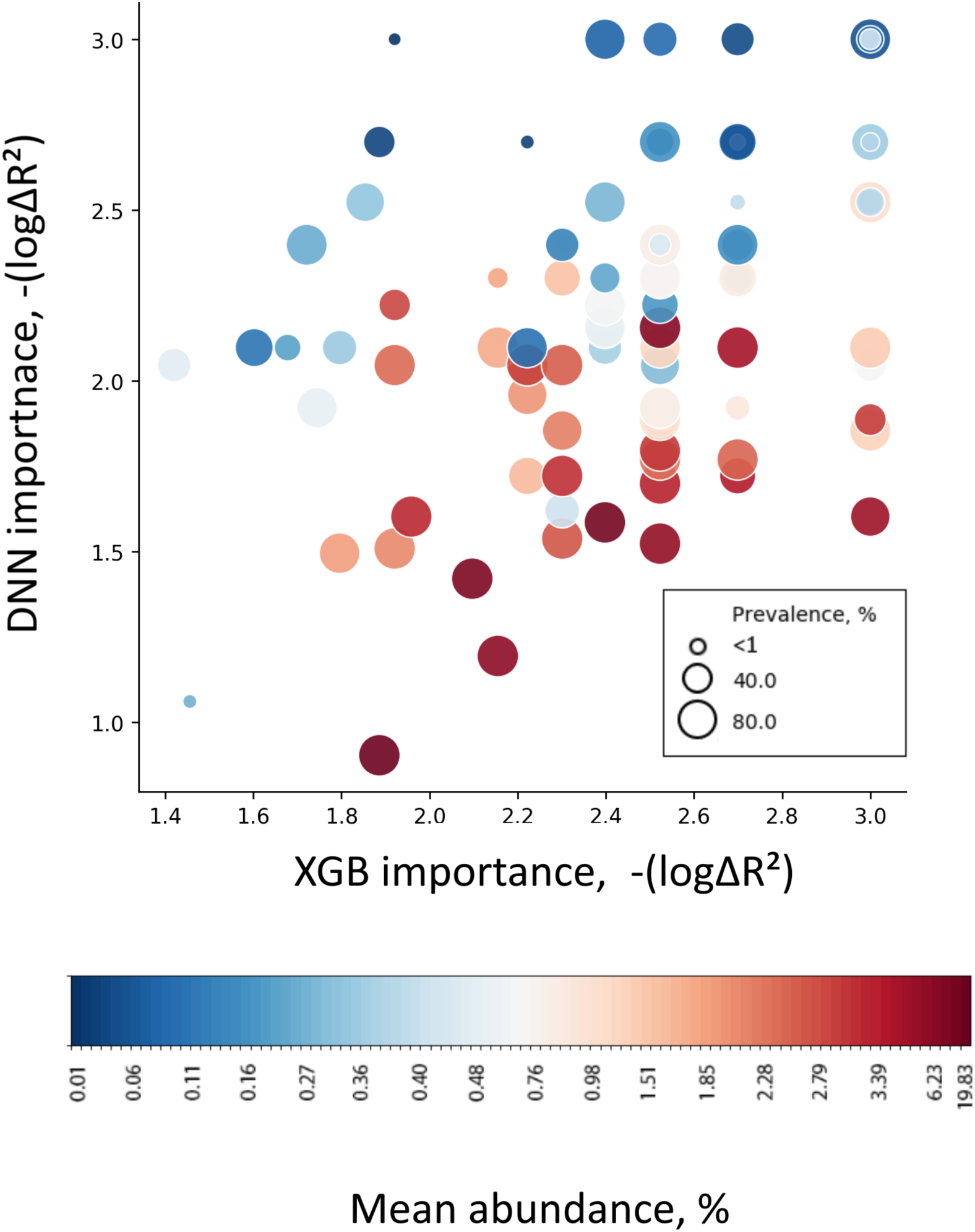
Bubble plot of 95 microbial taxa with PFI importance score >0.001 in both regression models (average among 5 folds). Bubble size stands for taxon prevalence (fraction of samples where the taxon is reliably detected), bubble color stands for taxon abundance (its average fraction in the communities where it was detected).

To characterize how these 95 features affect age prediction, we utilized the Accumulated Local Effects (ALE) approach (Figure 4). The ALE approach measures the response of a regressor to changes in specific taxa abundance. Each feature’s ALE was calculated using only the independent profiles where it can be reliably detected (abundance > 1e-5). Some microbes showed steadily increasing age prediction with increasing abundance (e.g. *[Eubacterium] hallii*); other microbes were on the opposite, inversely correlating with predicted age (e.g. *Bacteroides vulgatus*) (Figure 8). Interestingly, certain microbes that were previously identified as important by PFI showed little influence on predicted age (e.g. *[Eubacterium] rectale*) (Figure 8).

**Figure 8:**
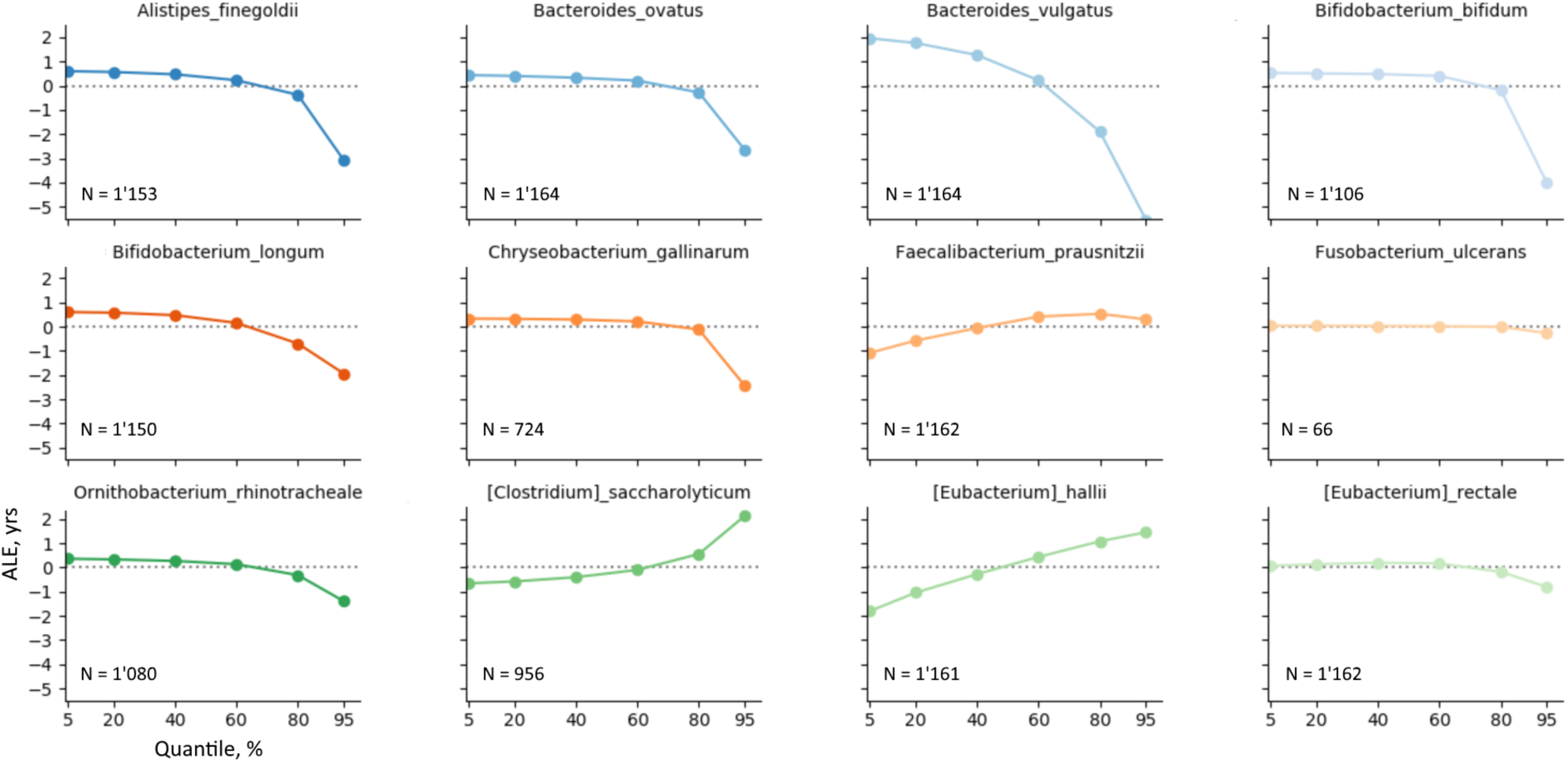
Twelve most important features’ effects on age prediction. Plots contain only 5-95% quantile segment due to extreme ALE values for extreme quantiles. N is the number of samples where a feature is reliably detected (abundance > 1e-05), total number of samples used is 1,165. More ALE plots are available in Supplementary Information.

Using ALEs, all features can be classified into seno-positive (monotonically increasing ALE plot), seno-negative (monotonically decreasing ALE plot), and more complex groups (not monotonic cases) (Figure 9). Among 95 features, only 39 displayed the average change in predicted age of more than 1 year within the 5%-95% quantile bracket. Among those, 21 were seno-negative, 15 seno-positive and 3 non-monotonic.

**Figure 9:**
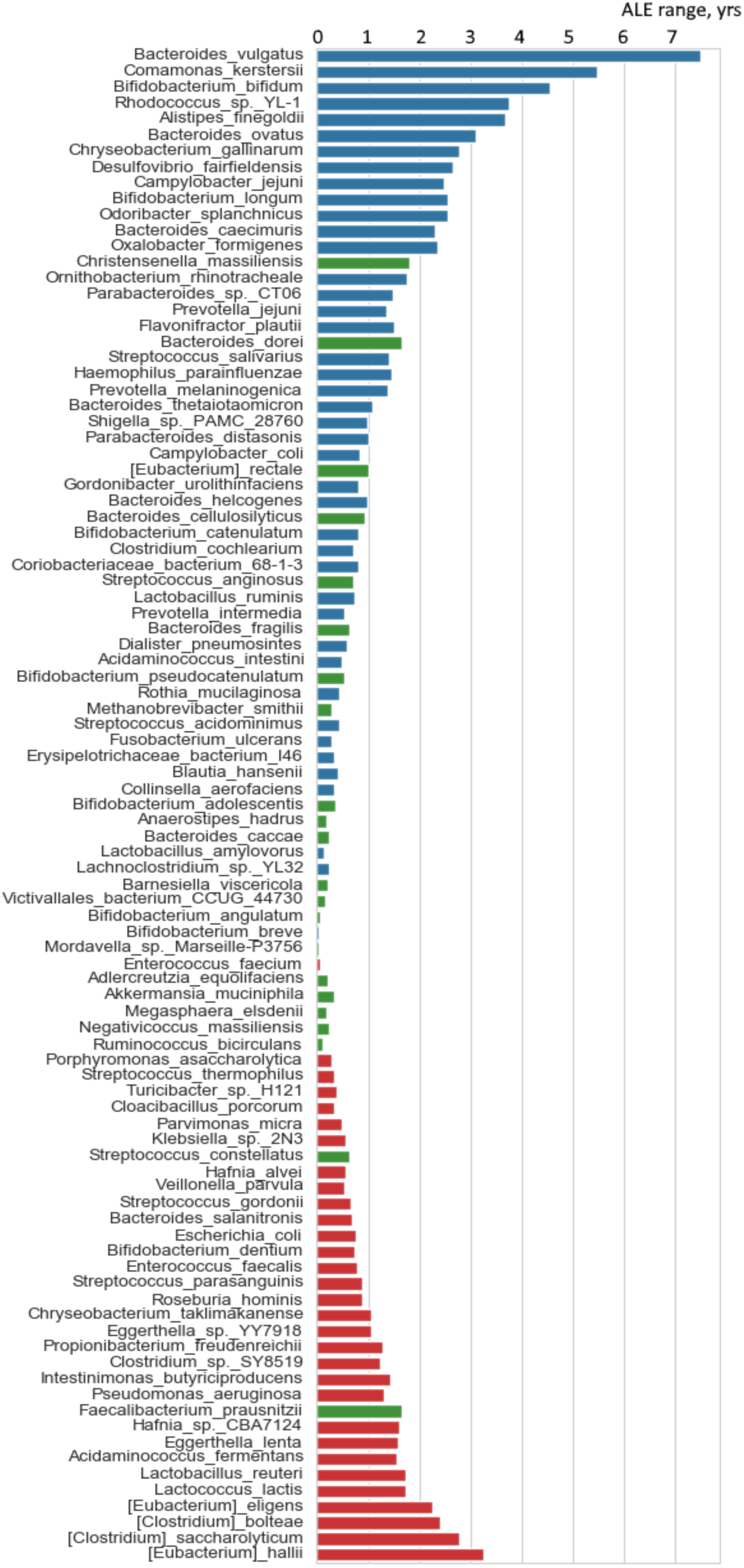
ALE range (maximum ALE minus minimum ALE within 5-95% abundance bracket) for 95 selected microbial features. Red are monotonically increasing ALEs, blue are monotonically decreasing ALEs, and green are non-monotonic ALEs. Only 39 taxa affect age prediction for more than 1 year within the specified abundance bracket.

### Age bracket prediction with DNN

While DNN and XGB regressors displayed acceptable accuracy when trained on full taxonomic profiles, decreasing the number of features down to 100 during training produces poorly performing models (MAE > 11 years). To estimate the predictive value of 95 and 39 marker taxa sets (Figure 9), we applied them to a much easier task of age bracket prediction. All donors were separated into three age groups: young (20-39 years, 32% of all donors), middle aged (40-59 years, 41% of all donors) and elderly (60-90 years, 27% of all donors). Underrepresented classes were oversampled (<u>see Methods</u>).

Within this setting, best performing DNN architectures show significantly higher accuracy than either random age group assignment (equiprobable or weighted). While the mean weighted F-score for random models do not exceed 38±1%, 95 marker set achieved the F-score of 67±4%. Downsizing this marker set using ALEs to 39 taxa reduced the score by 5% (to 62±3%). We have additionally compared the classifier constructed using the ALE-defined 39 intestinal marker set to classifiers built on relative abundances for 39 randomly selected taxa. Neither of 100 sets has produced a classifier as good as ALE-selected features (38±3%). (Figure 10)

**Figure 10:**
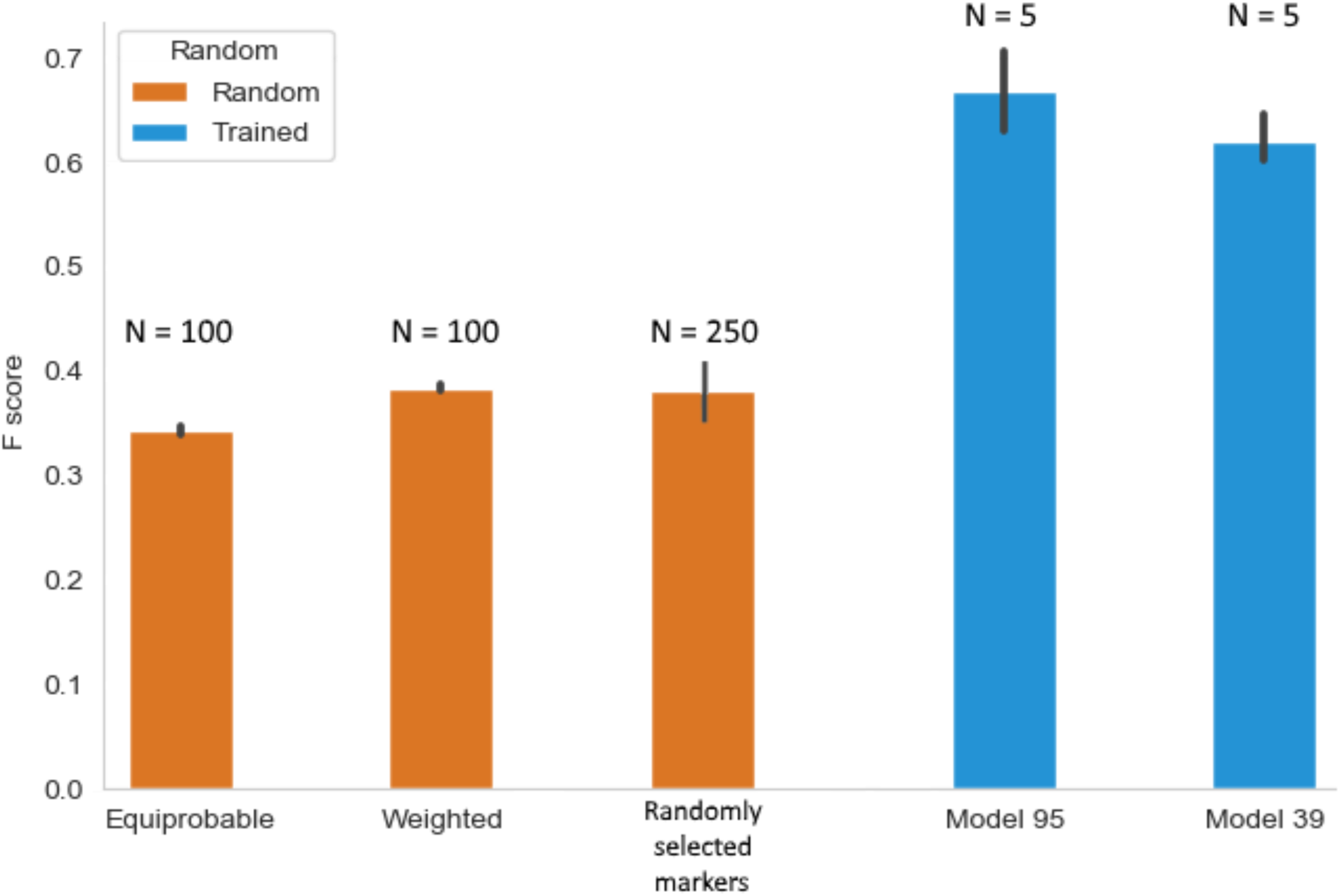
F-scores for four age bracket classifiers: three random models and two models built with 95 and 39 marker taxa. Equiprobable random classifier assigns a test sample to each age group (20-39, 40-59, 60-90 years) with ⅓ probability, weighted random classifier assigns samples with probabilities equal to the fraction of a class in the test sample. Models with randomly selected markers are built using 39 random taxa abundances as input. N stands for the number of cross validation folds.

### Host-based age prediction

While the DNN model is highly accurate, during its training all available samples were treated as independent due to data scarcity. By averaging the taxonomic profiles obtained from samples with a shared host we eliminated remaining data contamination. This reduced the total number of features to 1,165 entries. The host-based model was trained using the best performing DNN configuration as identified during sample-based training (Figure 2). This model was less accurate than a sample-based one: it reached MAE of only 8.56 years (Figure 11). However, the model still performed better than baseline age assignment (MAE = 12.47 years). Interestingly, the regressorprocesses feamle and male specimen with equal accuracy, and the predicted intestinal age positively correlates (r = 0.23) with BMI, which is in line with existing data on connections between BMI and biological age ^56^. However, this correlation is lower than the one between donor BMI and observed age: r = 0.3.

**Figure 11:**
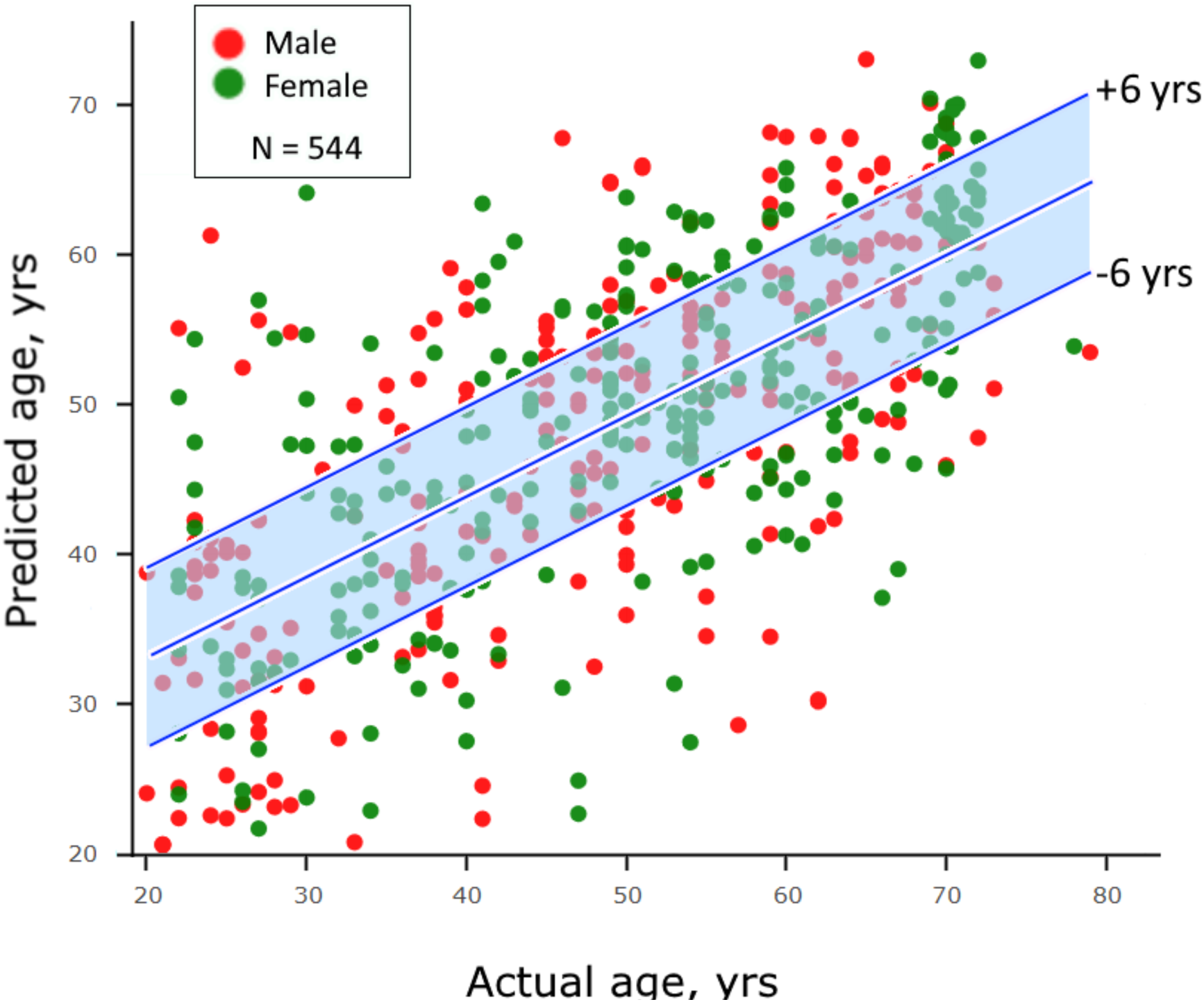
Age predictions derived from cross-validation of the host-based DNN model. Average MAE for best performing models in each of the 5 folds is 8.39 years, which is much lower than in the case of the sample-based approach (3.96 years). Blue area contains 52% of all predictions and corresponds to the trendline ±6 years.

## Discussion

To our best knowledge, we present the first method to predict human chronological age using gut microbiota abundance profiles. We compare two approaches to age prediction: regression and classification. We applied multiple methods to build a regressor that takes in profiles containing abundances for all 1,673 taxa reliably detected in at least 0.13% of samples, including random forest, support vector machine, elastic net, gradient boosting (XGB) and deep neural network (DNN). However, only the latter two models achieved the predictions better than random (Figure 5).

Due to data scarcity, we initially trained our models treating all samples as independent, while some of them belonged to the same host. To further demonstrate the applicability of the suggested method for age prediction, we trained a DNN model reducing the number of samples to only one per host. Not surprisingly, the resulting accuracy of the predictor was significantly lower (MAE = 8.56, Figure 11), yet above random. Such factors as study protocols and host country of residence (integrating geographic location, genotype and lifestyle) can be expected to affect taxonomic profiles.

Despite great performance of XGB (MAE = 4.69 years) and DNN models (MAE = 3.94 years), extracting biologically relevant information from them presents a major challenge. We implemented ALEs approach using DNN regressor as a reference and its 95 most important features to see how changes in abundance affect the predictions. ALE is a technique that theoretically surpasses PFI as it takes into account intrinsic interdependence of microbiological features. According to our ALE analysis, only 39/95 features could change the average predicted age by more than 1 year (Figure 9). Interestingly, reducing the number of features by 59% caused only a 5% drop in F-score for the age bracket classification task. This suggests that the ALEs technique succeeded in selecting only the most relevant microbial features.

Table 1 provides information for each bacterium in the 39 ALE-selected marker set of intestinal aging. Interestingly, while it contains both beneficial (e.g. *Bifidobacterium*) and pathogenic (e.g. *Pseudomonas aeruginosa*) microbes, seno-positive or seno-negative status is not determined by the nature of host-microbe interactions (Figure 12). For example, *Campylobacter jejuni* is known to cause campylobacteriosis – a foodborne diarrheal infection–yet it is seno-negative and can affect the average prediction age by more than 2 years (Figure 9) ^57^. On the other hand, both selected *Eubacterium* species are seno-positive and increase average predicted age by 1-3 years (Figure 9), despite having a generally beneficial effect on microbiota composition.

**Table 1:**
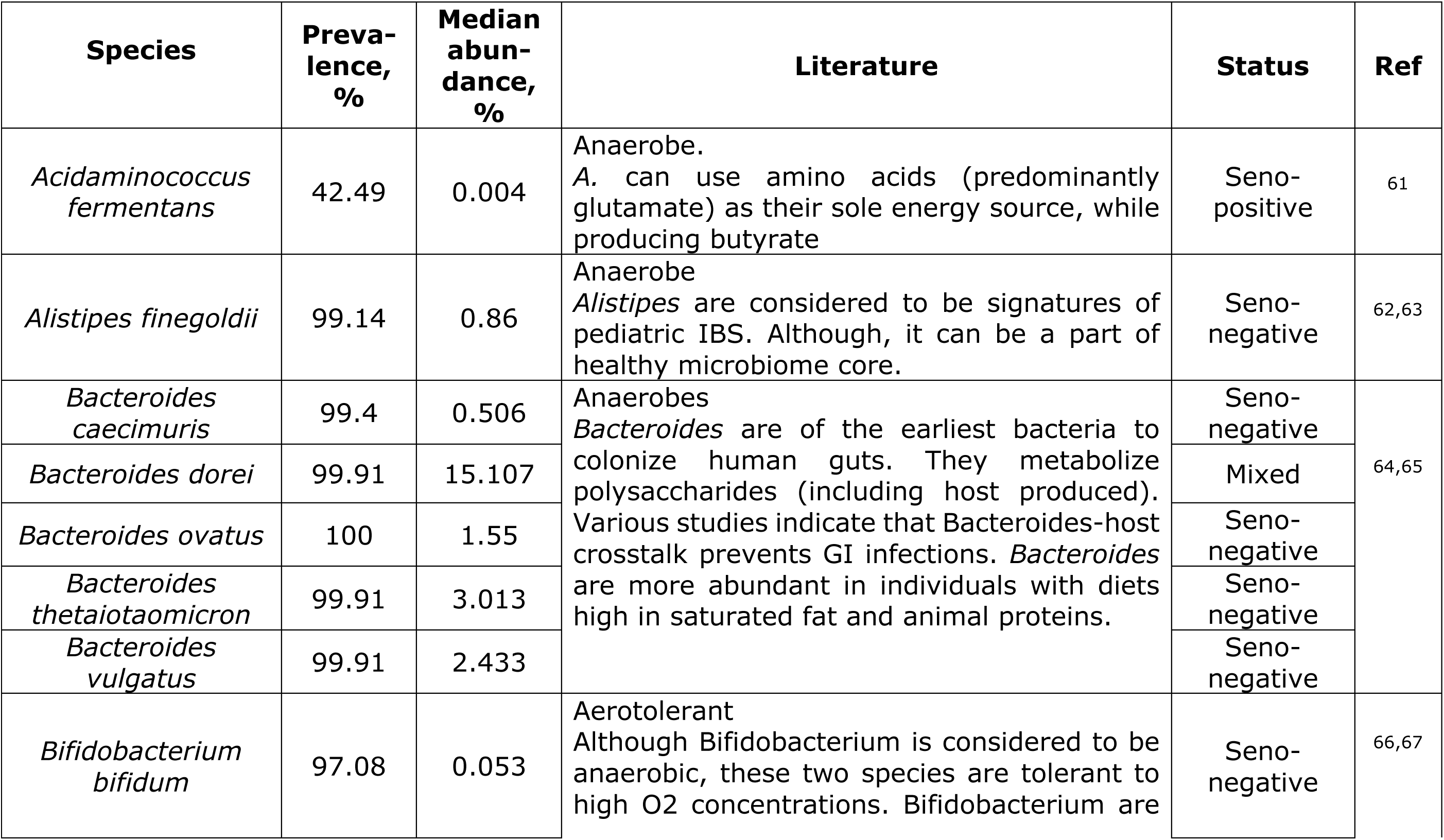
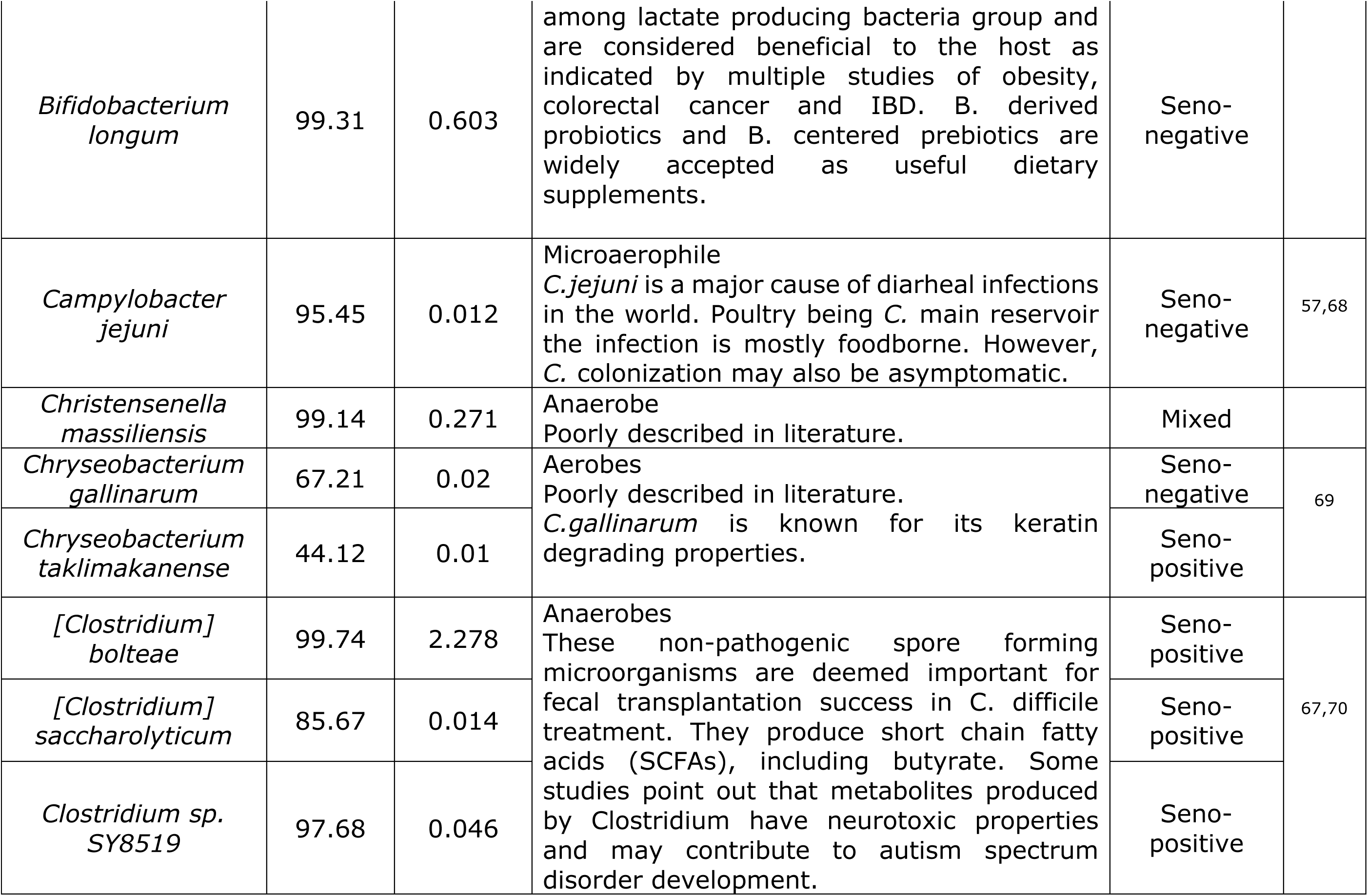
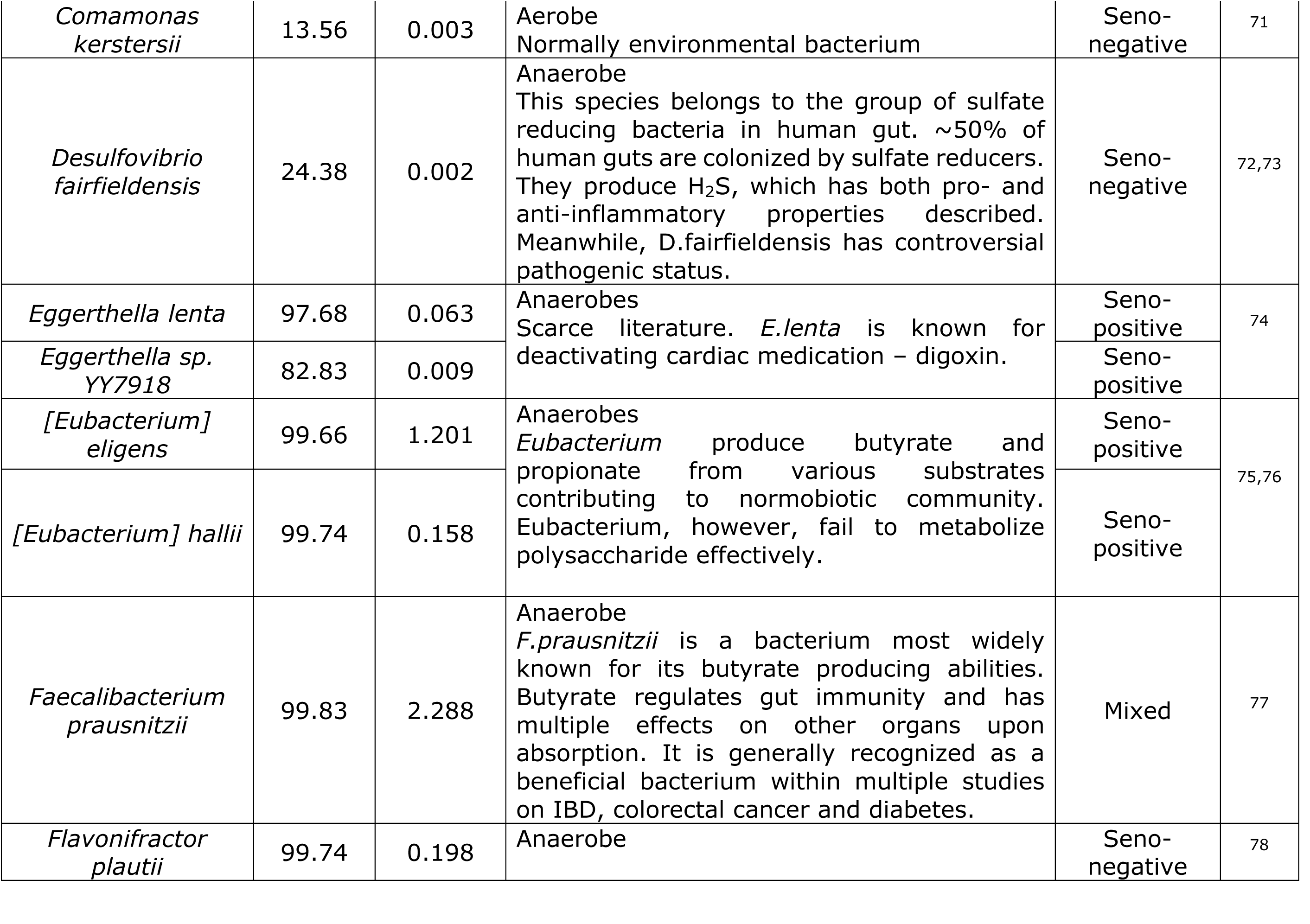
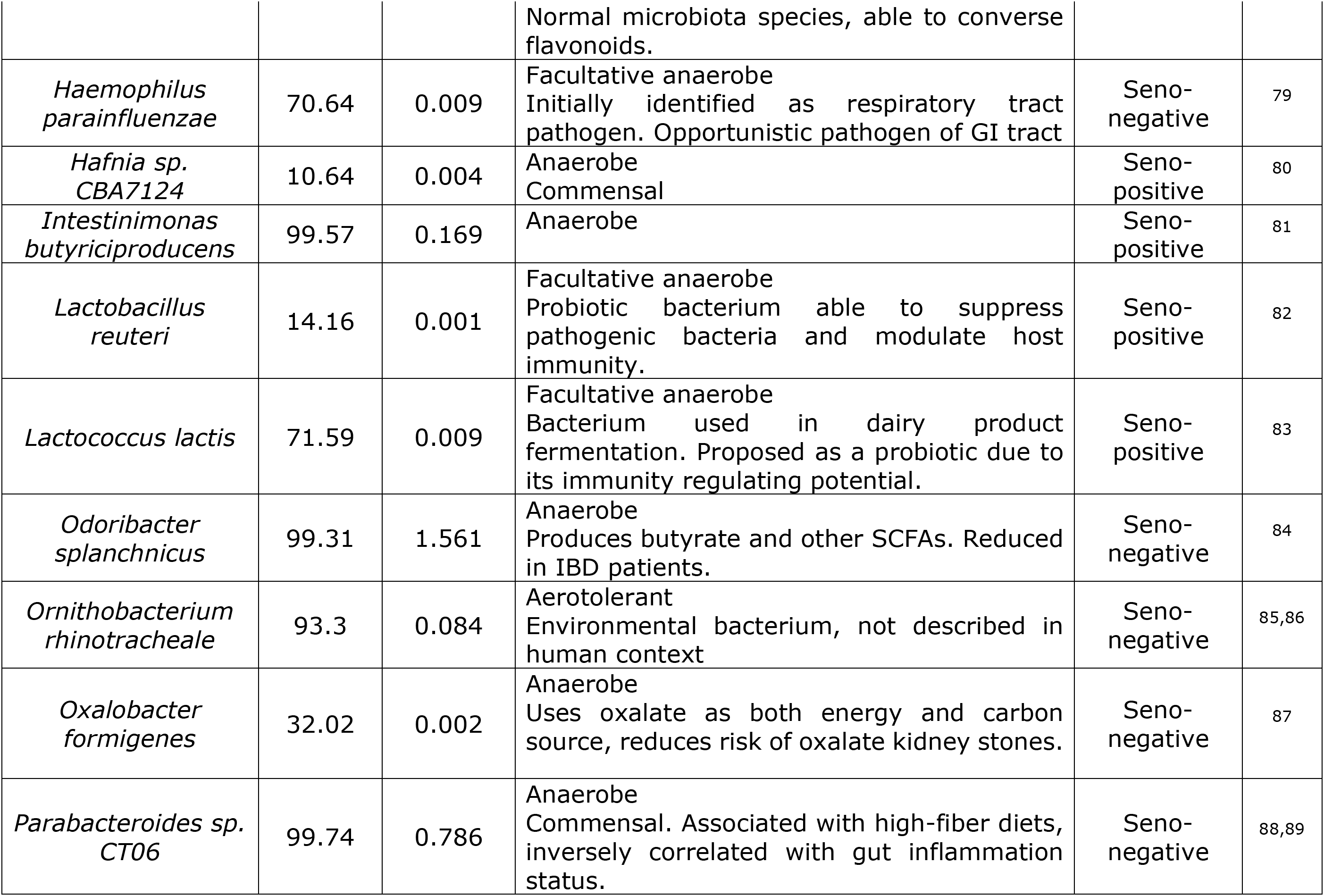
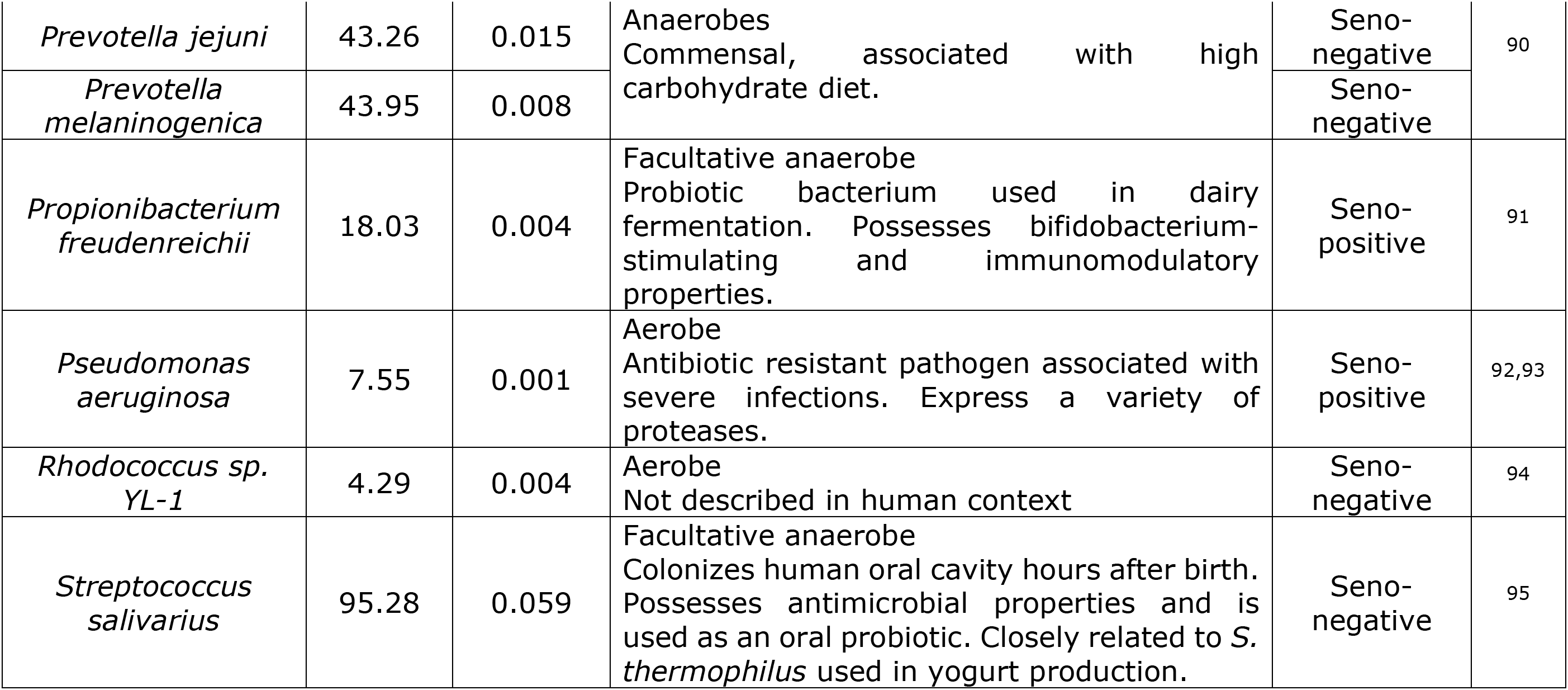
39 ALE-selected biomarkers for microbiological age prediction. Median abundance for a microbe is calculated excluding the samples where it is not detected. Total number of samples: 1,165

**Figure 12:**
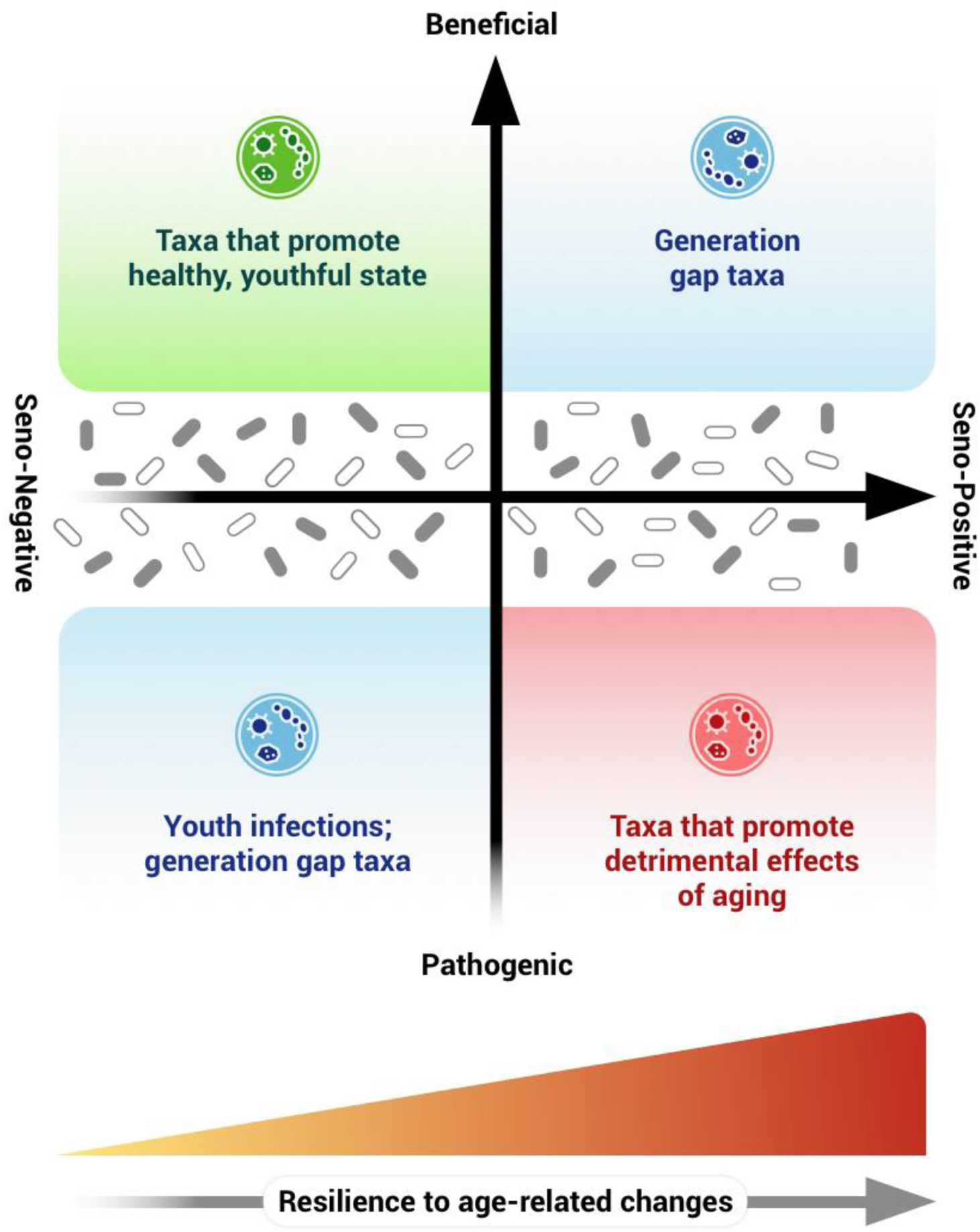
Seno-positive/-negative status of a species is not determined by its function within the gut community. While some pathogens are associated with increased age prediction (red quadrant) and some beneficial bacteria are correlated with lower age prediction (green quadrant), many other species are seno-negative pathogens and seno-positive normobiotic species (blue quadrants). Moreover, some taxa are scarcely described in literature, yet they have a pronounced effect on age prediction (grey area). In the text, we suggest hypothetical explanations as to why taxa might be occupying their respective positions in the plane above.

Although surprising at first glance, bacterial influence on age prediction is not determined by whether it is beneficial to the host or not. The proposed method of feature selection does not detect microbes that promote longevity or support useful functions of “youthful” microbiota. In the case of *C.jejuni*, campylobacteriosis affects mostly children. Moreover, exposure to *C.jejuni* can lead to asymptomatic colonization and immunity acquisition ^58^. Taken together, these facts can be used to put together a hypothetical explanation of *C.jejuni* being a seno-negative feature. Older individuals have a lower count of these bacteria, as they are more likely to carry the memory of previous *C.jejuni* exposure (either in their immune system or microbiota composition) and can effectively prevent its extensive colonization. Meanwhile, younger individuals have not yet tailored the means to oppose *C.jejuni* and let it multiply to a greater extent.

Beneficial bacteria that turn out to be seno-positive are, in fact, more intriguing than seno-negative pathogens. One possible explanation could be that these bacteria are more resilient in the context of increasingly detrimental cross-talk with the host. Another possible explanation questions the very concept of the microbiological aging clock. Since global dieting and lifestyle habits have significantly changed during the last century—increased sedentary time, sugar intake, processed foods consumption, etc.—any microbiota changes observed in the elderly may not indicate natural progression through age, but reflect generation-specific microflora parameters ^59^. In other words, any features identified today as associated with the youth may become the signature of the elderly in 50 years, provided global diet and lifestyle keeps changing. Depending on the extent of such microbiological “generation gap,” any future intestinal aging clock may need to be regularly updated to account for an ever-changing environmental context. Unfortunately, metagenomics is an extremely young branch of biology and there is little hope to learn what the microflora of the elderly looked like when they were young. However, studies of multiple generations of migrants can help us estimate the persistence of microbiota obtained in early years. One such study conducted on American immigrants of Asian origin indicates that first generation migrants start to lose gut diversity as soon as nine months after relocation. This loss is even more pronounced in the second generation ^60^. A first generation immigrant’s microflora memory may be the reason why their microbiota is different from that of their children. Studies in such a setting are extremely important to assess the hyper-parameters driving human microbiota progression.

Another interesting aspect of the ALE-based feature selection is that some features are very poorly described within human microbiota context—environmental bacteria, —yet have great influence over age prediction. More knowledge on such microorganisms (e.g. *Ornithobacterium rhinotracheale*) may provide useful insights into the functions of human microbiota.

## Conclusion

We demonstrated the feasibility of age prediction by application of machine learning approaches to taxonomic microflora profiles. Our most accurate DNN regressor achieved the MAE of 3.94 years. This performance is comparable with the 1.9 MAE of the PhotoAgeClock, 2.7 of the state of art methylation aging clock, 7.8 MAE transcriptomic aging clock and 5.5 MAE of the hematological aging clock published previously. We also developed a method for microbiological feature selection and annotation. It combines two-fold feature importance assessment using PFI and ALE approaches upon training a DNN. This technique allows both selecting the most relevant features as biomarkers and quantifying their influence on the target variable, i.e. age. Using this method, we identified 95 and 39 prokaryote taxa as the biomarkers of intestinal aging. Despite the reduced predictive power of this set when compared to the whole taxonomic profiles, it let us to assign individuals to three age groups (young, middle aged and old) 86% more accurately than random classification (0.71 versus 0.34 F-score).

The identified biomarkers include species whose abundance is positively or negatively correlated with predicted age. These species may be further investigated deeply by the community to improve our understanding of human aging and its relationship with the gut microbiome.

## List of abbreviations

ALE: Accumulated local effect
BMU: Best matching unit
DFS: Deep feature selection
DNN: Deep neural network
IBD: Inflammatory bowel diseases
IBS: Irritable bowel syndrome
LE: Local effect
MAE: Mean absolute error
OUT: Operational taxonomic unit
PFI: Permutation feature importance
SCFA: Short chain fatty acids
SOM: Self-organizing maps
WGS: Whole genome shotgun [sequencing]
XGB: Gradient boosting (XGBoost Python implementation)

## Acknowledgments

This publication contains free vector art created by <u>Alice Noir</u> from the Noun Project and by <u>eucalyp</u> from Flaticon. VNG is supported by NIH grants.

